# A simple MiMIC based approach for tagging endogenous genes to visualise live transcription *in vivo*

**DOI:** 10.1101/2024.08.29.610339

**Authors:** Lauren Forbes Beadle, Catherine Sutcliffe, Hilary L. Ashe

## Abstract

Live imaging of transcription in the *Drosophila* embryo using the MS2 or PP7 systems is transforming our understanding of transcriptional regulation. However, insertion of MS2/PP7 stem loops into endogenous genes requires laborious CRISPR genome editing. Here we exploit the previously described Minos-mediated integration cassette (MiMIC) transposon system in *Drosophila* to establish a method for simply and rapidly inserting MS2/PP7 cassettes into any of the thousands of genes carrying a MiMIC insertion. In addition to generating a variety of stem loop donor fly stocks, we have made new stocks expressing the complementary coat proteins fused to different fluorescent proteins. We show the utility of this MiMIC-based approach by MS2/PP7 tagging and live imaging transcription of endogenous genes and the long non-coding RNA, *roX1*, in the embryo. We also present live transcription data from larval brains, the wing disc and ovary, thereby extending the tissues that can be studied using the MS2/PP7 system. Overall, this first high throughput method for tagging mRNAs in *Drosophila* will facilitate the study of transcription dynamics of thousands of endogenous genes in a range of *Drosophila* tissues.

## Introduction

Gene expression underlies every aspect of an organism’s development and homeostasis. The expression of genes at particular times or in specific cells is exquisitely controlled to ensure correct development. Transcriptional control is mediated at the level of DNA accessibility, transcription factor recruitment and RNA polymerase II initiation, elongation and termination (Cramer, 2019; Haberle and Stark, 2018; Hager et al., 2009; van Steensel and Furlong, 2019). Many tools have been developed to visualise and measure mRNAs within cells, including *in situ* hybridisation-based methods and genomics approaches such as RNAseq (Pichon et al., 2018; Stark et al., 2019). In addition, the development of live imaging techniques allows transcription to be visualised within cells at active gene loci (Ferraro et al., 2016; Pichon et al., 2018).

The bacteriophage derived RNA stem loops, MS2 and PP7, are used to study nascent transcription and mRNA localisation in living cells (Bertrand et al., 1998; Chubb et al., 2006; Larson et al., 2011). Multiple copies of these loops are inserted into the gene of interest using CRISPR or transgenesis. When the gene is transcribed by RNA Polymerase II, the RNA stem loop structures are formed and bound by a co-expressed fluorescently tagged coat protein, MS2 coat protein (MCP) or PP7 coat protein (PCP) (Larson et al., 2011). This allows the fluorescent signal from each transcription site to be visualised and quantitated over time, with the signal proportional to the number of RNA Polymerase II molecules transcribing the gene. Live imaging has revealed bursts of transcriptional activity that can be described by a two state promoter model, where the promoter switches between active (ON) and inactive (OFF) states (Golding et al., 2005; Lionnet and Singer, 2012). Control of the promoter states is tuned by transcription factor inputs and is important for proper developmental processes (Garcia et al., 2020).

As a premier model organism, *Drosophila* has many genetic tools available to the scientific community. The collection of MiMIC insertions are stocks containing transposable elements with inverted ΦC31 recombinase target sites which are spread across the *Drosophila* genome and can be used for gene tagging via cassette exchange (Nagarkar-Jaiswal et al., 2015a; Nagarkar-Jaiswal et al., 2015b; Venken et al., 2011). Currently there are 7451 insertion stocks available at the Bloomington stock centre associated with 4367 distinct genes, which can be targeted using this technology. MiMIC insertions have an advantage over other types of genomic inserted elements (pBac and P elements) as they have no sequence bias and can insert in any genomic location. Therefore, the insertion collection targets noncoding regions of genes, including the UTRs and intronic sequences (Venken et al., 2011). The MiMIC insertions have been genetically manipulated to tag genes with various sequences to study protein expression and localisation (Nagarkar-Jaiswal et al., 2015a; Nagarkar-Jaiswal et al., 2015b). This is achieved by recombination mediated cassette exchange (RMCE) at the MiMIC sites within introns, without the need for microinjection. As a result, this approach enables gene tagging, as crossing two fly lines to genetically manipulate a gene region is much easier than having to design cloning strategies and perform microinjections to achieve the same outcome.

Here, we describe the generation of a new set of fly stocks that can be used to insert a variety of MS2/PP7 stem loops into any of the thousands of genes within the existing MiMIC library. We have also made five new coat protein-fluorescent protein stocks that improve detection options. We validate the approach by inserting different MS2/PP7 sequences into protein coding genes and the *roX1* long non coding RNA (lncRNA) to study their transcription dynamics in living embryos. We have also used our new lines to extend live transcription imaging to other tissues during development, including the ovary, wing disc and larval brain. Overall, as the MiMIC-based approach relies on a simple crossing scheme, the new stocks we have generated will greatly facilitate the study of transcription of endogenous genes in living cells *in vivo*.

## Results

### A new MS2/PP7 toolkit for MiMIC insertion and transcription live imaging

Live imaging of nascent transcription requires the insertion of an array of stem loop hairpin sequences (MS2 or PP7) into the gene of interest. These hairpins are bound by MCP or PCP respectively, expressed as a fluorescent protein fusion (Fig. 1Ai). Transcription sites (TSs), which are visible as bright fluorescent puncta within each nucleus, can be tracked over developmental time to reveal transcription dynamics (Fig. 1Aii).

**Figure 1.**
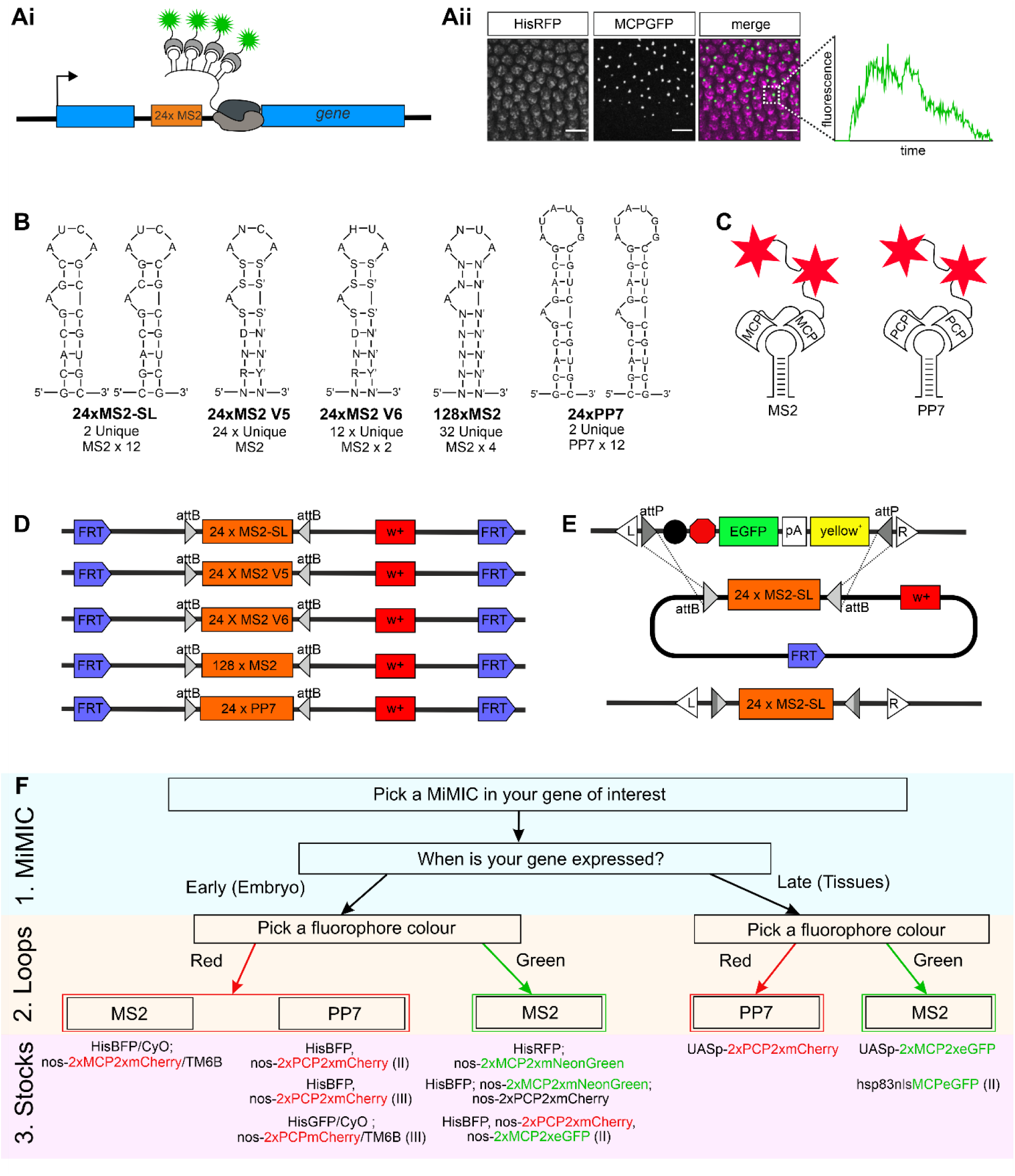
Overview of the MiMIC insertion strategy for MS2/PP7 sequences. (Ai) Schematic shows 24xMS2 loops (orange box) inserted into a gene intron. Following transcription, fluorescent coat proteins (green stars) bind to the stem-loops in the RNA. (Aii) Still from a live imaging movie showing TSs, visualised as bright fluorescent puncta of MCPGFP bound to the stem loops, within the nuclei marked by HisRFP. Fluorescent intensity can be tracked over time as shown for one TS in a nucleus (inset). Scale bar = 10µm. (B) Generalised nucleotide sequences of the MS2 and PP7 loop versions used in this study. (C) Schematics of MCP bound to a MS2 loop and PCP bound to a PP7 loop. In each case a tandem dimer coat protein is fused to two fluorescent proteins (red stars). (D) Schematics of the plasmids injected into flies to create the loop donor fly stocks. (E) Scheme for insertion of the loop donor cassette into the MiMIC containing locus. FLP/FRT mediated removal of the cassette is followed by RMCE at the attB/attP sites within the donor and the target site, mediated by germline expression of the ΦC31 integrase (top). This results in integration of the loops into the MiMIC site within the gene (bottom). (F) A flowchart showing the three genetic components required for visualisation of live transcription *in vivo*: 1) the MiMIC insertion, 2) the loop donor constructs and 3) the corresponding coat protein fly stocks generated or used in this study.

We generated several fly lines that carry insertions with different versions of the MS2 or PP7 loops (Fig. 1B), including 24xMS2-SL, 24xMS2V5, 24xMS2V6, 128xMS2, 24xPP7 (Bertrand et al., 1998; Fukaya et al., 2016; Tantale et al., 2016; Tutucci et al., 2017). The version of the MS2 loops that were optimised for stable expression in bacteria (MS2-SL) has been used in many live imaging studies in the *Drosophila* embryo (Fukaya et al., 2016; Garcia et al., 2013; Hoppe et al., 2020; Lammers et al., 2020; Lim et al., 2018). MS2-SL consists of twelve pairs of repetitive sequences that form stem loop structures with short linker sequences in between. MS2V5 is an array of 24 non-repetitive stem loop sequences (Tutucci et al., 2017). The MS2V6 variant contains a U instead of a C at a crucial residue in the coat protein binding site that makes the interaction weaker and the loop-coat protein interaction less stable (Tutucci et al., 2017). A strong coat protein-loop interaction has been shown to interfere with mRNA degradation in yeast cells when the loops are in the 3’ UTR (Garcia and Parker, 2015). Although the effect of stem loops on mRNA degradation has not been addressed in *Drosophila*, the V6 version may be useful for studies in which correct mRNA degradation of the target is important. The 128xMS2 loop variant increases the number of loops, which improves the detection and signal to noise ratio, allowing weak TSs and single mRNAs to be detected (Dufourt et al., 2021; Tantale et al., 2016; Vinter et al., 2021). 24xPP7 is a different stem loop structure that can be utilised in combination with MS2 to allow dual colour imaging of each allele of the same gene or two different RNAs.

Once the stem loops have been transcribed by RNA Pol II, two MCP or PCP proteins bind to each loop (Fig. 1C). Expressing the coat proteins as a tandem dimer improves complete occupancy at all stem loops and quantitation of the fluorescent signals (Wu et al., 2012). Therefore, we have generated new stocks with MCP/PCP tandem dimers fused to two fluorescent proteins to maintain the ratio of two fluorescent proteins for each stem loop (Fig. 1C). MCP/PCP have been fused to different fluorescent proteins and the new lines typically also carry HisBFP (listed in Fig. 1F) to simultaneously visualise the nuclei during imaging.

We utilised the previously described genetic strategy to design these lines so they can be used with RMCE to genetically insert the loop array sequences into MiMIC containing genes (Nagarkar-Jaiswal et al., 2015a). The MS2/PP7 loop array cassettes were cloned between inverted attB sites (Fig. 1D), which lie upstream of a *mini-white* (*w^+^*) marker that is used for selection and to follow inheritance of the cassette. The entire cassette is flanked by FRT sites, allowing FLP mediated excision. As our constructs are to be used for visualising nascent transcription, these donor lines do not need splice acceptor or donor sites, or need to be in frame, unlike previous studies that used MiMIC insertions for protein localisation (Nagarkar-Jaiswal et al., 2015a). Using a crossing scheme and heat shock, RMCE inserts the repetitive loop sequences into the gene of interest with the action of germline expressed *vasa* ΦC31 integrase at the inverted attP sites (Fig. 1E). The three genetic components required to visualise transcription are shown in Fig. 1F. Firstly a MiMIC insertion in the gene of interest is required, secondly the type of loop is selected and thirdly the fluorescent coat protein stock is chosen.

### Overview of the protocol for inserting loops into MiMIC containing genes

A simple three step method is used to insert the loop cassette sequence into the MiMIC insertion site of a gene of interest (Fig. 2A). MiMIC insertions are readily available from the Bloomington stock centre and all the stocks described here will also be made freely available. Step 1 is the selection of the loop type and the gene of interest containing a MiMIC insertion. In this example, we inserted 24xPP7 loops into the second intron of the *pxb* gene using the Mi04897 MiMIC insertion by crossing virgin females from the PP7 stem loop donor flies to males with the MiMIC insertion. At step 2, the F1 embryos and larvae are heat-shocked which liberates the loop cassette. Activation of the heat shock FLP recombinase results in mosaic eyed progeny which should be selected for in the emerging adults. The loop cassette can then insert into the region between the inverted attP sites of the MiMIC locus via RMCE, which results in loss of the MiMIC’s *yellow*+ (*y*+) marker in the next generation.

**Figure 2.**
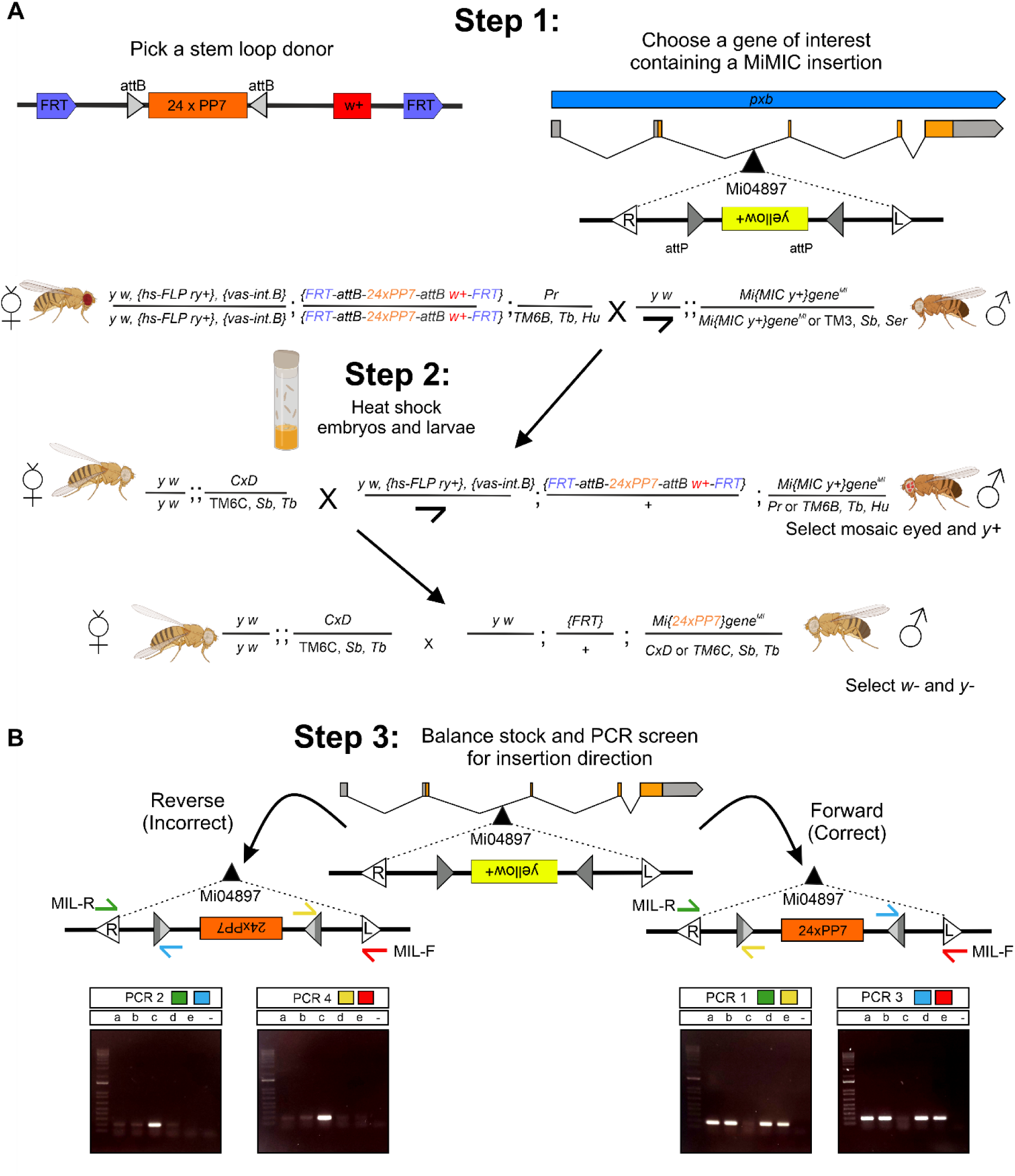
Step by step protocol for inserting loops into MiMIC containing genes. (A) Crossing scheme to target a MiMIC containing gene on the third chromosome (*pxb*) with 24xPP7 loops. The *pxb* gene contains an intronic MiMIC insertion (Mi04897). The Mi04897 insertion is in the opposite orientation to *pxb* and is marked by the *yellow+* marker and flanked by inverted attP sites (grey triangles). The selection of markers as indicated will result in flies with loops inserted into the MiMIC. (B) PCR screening of 24xPP7 loops inserted into *pxb* at Mi04897. The opposite direction relative to the gene is shown by a band in PCR2 (green and blue primers) and PCR 4 (yellow and red primers) and the absence of a band in PCR1 (green and yellow primers) and PCR 3 (blue and red primers). The same direction relative to the gene is confirmed by a band in PCR1 (green and yellow primers) and PCR 3 (blue and red primers) and the absence of a band in PCR2 (green and blue primers) and PCR 4 (yellow and red primers). Samples a, b, d and e here are in the correct forward orientation. MIL-F (red primer) and MIL-R (green primer) are common to all MiMIC inserts and the internal yellow and blue primers are stem-loop specific.

After the stock has been established in step 3, the male parent is sacrificed for PCR screening and/or sequencing to determine the orientation of the stem loop insert (Fig. 2B). All *yellow white* (*yw*) males will have an insertion event but as the cassette can insert in either direction we recommend that at least 5-10 *yw* males are screened to ensure an insertion in the correct orientation. Combinations of PCR primers overlapping the Minos L and R sequences and near the attL/R sites within the stem loop insert can be used to determine the orientation of the loops. In the example shown here, sample c with a PCR product for primer sets PCR 2 and PCR4 had the PP7 loops inserted backwards relative to the *pxb* gene, whereas samples a, b, d and e with a PCR product for primer sets PCR1 and PCR3 had the insertion in the same orientation as the gene (Fig. 2B). Dual testing of the direction of the insert will give confidence that the loops are inserted in the correct orientation to be bound by the fluorescent coat proteins for visualisation (Valegård et al., 1994). Additional sequencing of the PCR product can be used to confirm the correct direction of the loops.

In summary, this crossing scheme is easy and efficient so that lines with integrated MS2/PP7 sequences integrated into the gene of interest can be obtained in as little as a month. The crossing schemes for insertion of a loop cassette into the X and second chromosome are shown in Fig. S1. Considerations for targeting a gene on the X chromosome are discussed in the Methods. Once the desired stock has been generated it can be crossed to fluorescent coat protein lines to visualise live transcription *in vivo*. We find no evidence that the insertion of these loops interferes with the endogenous expression pattern of any of the genes we have investigated to date, including those in this study (Fig. S2).

### Transcription live imaging of both *pxb* alleles in the *Drosophila* embryo

To show the utility of the MiMIC tagging live imaging system, we used the new lines we generated that contain either 24xMS2V6 loops or 24xPP7 loops in the *pxb* locus (Fig. 2) to visualise nascent transcription in the early *Drosophila* embryo. HisRFP; nos-2xMCP2xmNeonGreen females lay embryos with these proteins maternally loaded. After crossing these females to *pxb*-24xMS2V6 males, live imaging the resulting embryos will detect RFP labelled nuclei and fluorescent *pxb* TS puncta labelled with mNeonGreen (Fig. 3Ai). Live imaging embryos towards the end of nuclear cycle 14 allows visualisation of the TS puncta within the nuclei of cells within the *pxb* expression domain (Fig. 3Aii, Movie 1). The endogenous pattern of *pxb* is striped across the embryo at this stage (Inaki et al., 2002); only two of the anterior stripes of the expression domain are shown in the movie still (Fig. 3Aii), as highlighted by the yellow region on the schematic (Fig. 3Ai). The TSs detected are consistent with the endogenous expression of *pxb* within this region (Fig. S2).

**Figure 3.**
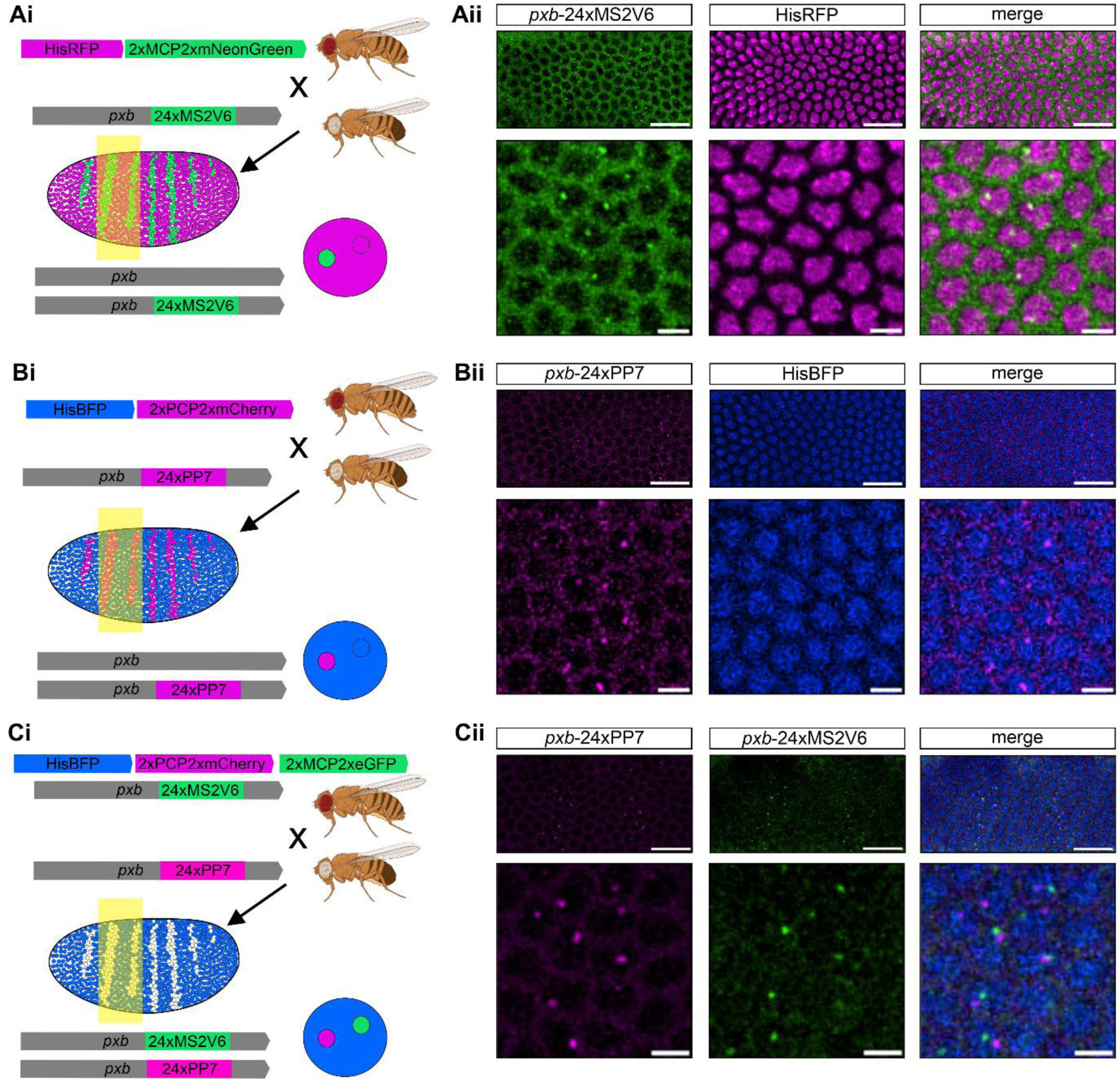
Transcription live imaging of *pxb* tagged with MS2 and PP7 stem loops in embryos. (Ai) Schematic showing the *pxb* gene locus with the 24xMS2V6 insertion. Males with the 24xMS2V6 loops inserted into *pxb* are crossed to females expressing HisRFP and 2xMCP2xmNeonGreen under the control of the *nos* promoter. Embryos have one of the two *pxb* alleles visible as a mNeonGreen puncta within the HisRFP nucleus, the imaging region is shown in yellow. (Aii) Top: Still images from a timelapse movie of *pxb-*24xMS2V6 transcription sites in two anterior stripes in the nc14 embryo. Scale bar is 20µm. Bottom: Higher magnification image from nuclei with the stripe. Nuclei are marked in magenta and *pxb-*MS2 TSs are in green. Scale bar is 5µm. (Bi) As in (Ai) but with 24xPP7 loops inserted into *pxb* and females expressing HisBFP and 2xPCP2xmCherry. (Bii) Still from a movie of an embryo expressing HisBFP and 2xPCP2xmCherry marking the nuclei in blue and the *pxb-*PP7 TSs in magenta. (Ci) Schematic showing *pxb* labelled to visualise each allele with a different loop type (MS2 or PP7). (Cii) Dual imaging of these embryos detects transcription of both alleles. *Pxb-*PP7 TSs are in magenta, *pxb-*MS2 TSs are in green, nuclei are blue in the merge. See also Movies 1-3.

Next, we tested the *pxb*-24xPP7 insertion by crossing males to females expressing HisBFP and 2xPCP2xmCherry (Fig. 3Bi). Imaging detected mCherry fluorescent *pxb* TS foci within BFP labelled nuclei. The *pxb*-24xPP7 embryos show the same expression pattern in the anterior stripes as described for the *pxb*-24xMS2 embryos (Fig. 3Bii, Movie 2). As two different types of loops were inserted in the *pxb* locus, we crossed the lines together to visualise both alleles at the same time (Fig. 3Ci, Cii, Movie 3). Analysis of this type of data can be used to infer transcriptional burst parameters and study the degree of coordination of transcription between both alleles.

### Live imaging of transcription of the BMP target gene *Race* and *roX1* lncRNA

We next targeted the other major chromosomes in *Drosophila* and inserted loops into genes on the X and second chromosomes. For the second chromosome, we chose the Dpp target gene *Race* (also known as *Ance*) (Ashe and Levine, 1999; Tatei et al., 1995). *Race* contains a MiMIC insertion Mi05748 within the first coding intron (Fig. 4A). We used this MiMIC insertion to insert 24xPP7 loops into the endogenous locus as described in Fig. 2 using the crossing scheme in Fig. S1B. For the X chromosome, we used Mi01457 to insert 24xPP7 loops within the *RNA on the X 1* (*roX1*) lncRNA, which plays a critical role in dosage compensation in males (Samata and Akhtar, 2018) (Fig. 4B).

**Figure 4.**
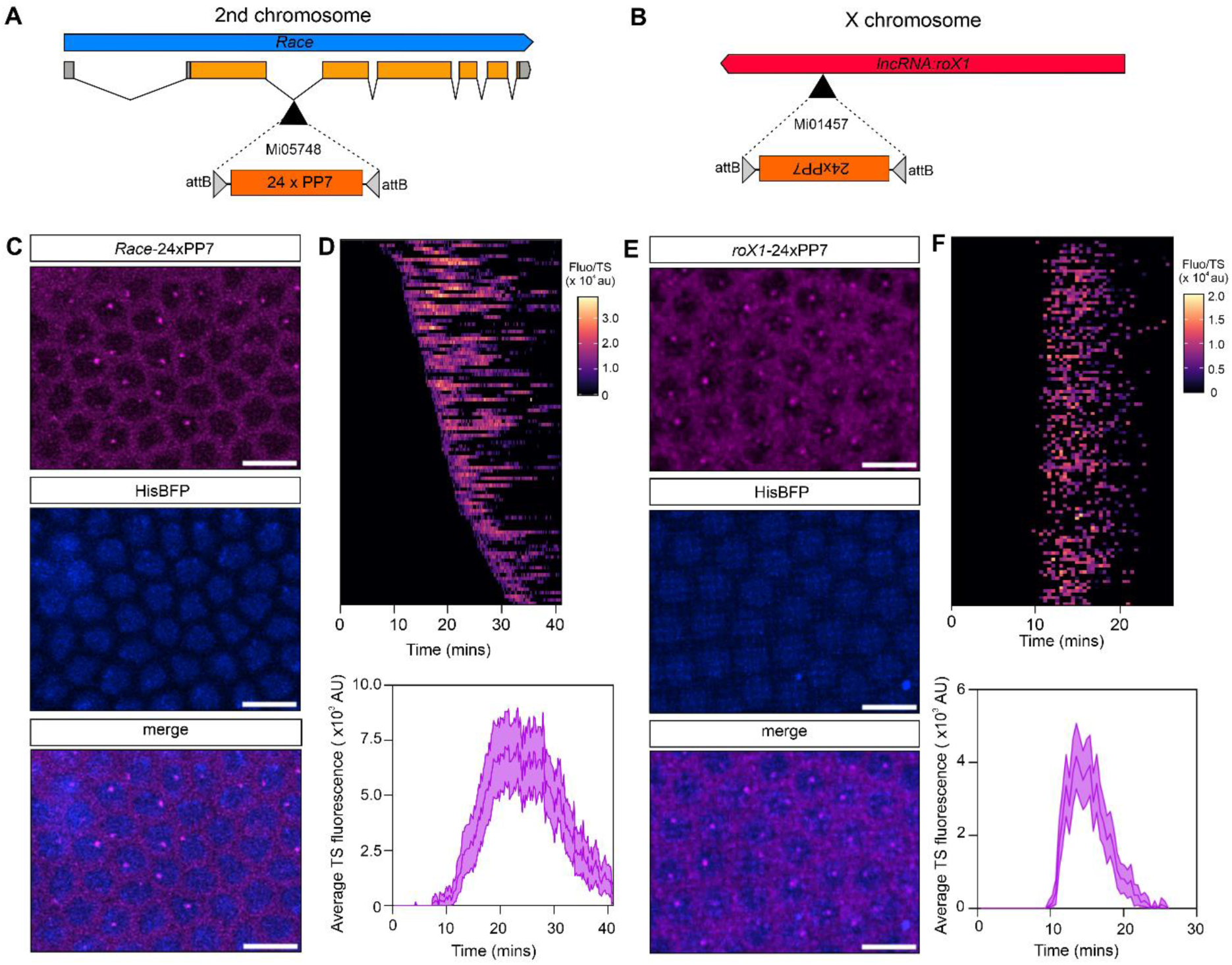
Live imaging of *Race* and *roX1* transcription. (A, B) 24xPP7 loops were inserted into (A) the second chromosome gene *Race* at Mi05748 and (B) the X chromosome lncRNA *roX1* at Mi01457. (C) Stills from a live imaging movie of an embryo with *Race*-24xPP7 and maternally expressed HisBFP and 2xPCP2xmCherry under the control of the *nos* promoter, showing nascent transcription sites within the *Race* expression domain. (D) Top: Heatmap of individual *Race* transcription site traces measured during nc14. Bottom: Graph showing mean TS fluorescence intensity of *Race* in a nc14 embryo. (E and F) As in (C) and (D) but the data are shown for *roX1*-24xPP7. Mean ± CI for 72 nuclei (*Race)* and 115 nuclei (*roX1*). Scale bar is 20µm.

Females expressing HisBFP and 2xPCP2xmCherry were crossed to *Race*-24xPP7 males and the resulting embryos were imaged during nc14 (Movie 4). A representative image of the anterior dorsal view of the embryo shows nuclei transcribing *Race* during this time (Fig. 4C). *Race* TSs were tracked and the single nuclear transcriptional fluorescent traces show stochastic onset in nc14 (Fig. 4D). Calculating the mean TS fluorescence shows that transcription peaks ∼20 min into nc14 (Fig. 4D).

HisBFP; 2xPCP2xmCherry females were crossed to *roX1*-PP7 males and embryos collected to detect *roX1* transcription across early development. Ubiquitous transcription of this lncRNA was observed during nc14 (Fig. 4E). Transcription initiates synchronously and was detected in a short time period from approximately 10-25 minutes into nc14 (Fig. 4F and Movie 5). Together, these experiments and those shown for *pxb* validate that we can use the MiMIC approach to target MS2/PP7 loops into the three major chromosomes in *Drosophila.* This has allowed us to visualise and track transcriptional activity of not just mRNAs, but also a lncRNA, which opens up new possibilities for studying non coding RNA transcriptional dynamics.

### Live imaging of transcription in the ovary, wing disc and larval brain

We next addressed whether the new stem loop lines we have generated could be used to visualise nascent transcription in other tissues later in development. To this end, we constructed new fly stocks for expression of the fluorescent coat proteins. We first tested the GAL4-UAS system (Brand and Perrimon, 1993) and made UASt (Brand and Perrimon, 1993) and UASp (Rørth, 1998) versions of 2xMCP2xeGFP and 2xPCP2xmCherry. However, we found that expressing UASt-2xMCP2xeGFP or UASt-2xPCP2xmCherry in somatic tissues, such as the wing disc, with GAL4 drivers resulted in very high expression levels that caused accumulation of excess coat protein fusions in intense foci. As a result, TSs were undetectable (data not shown). Therefore, we next tried using UASp to drive expression in the somatic tissues using the same drivers, as UASp transgenes have a much lower level of expression in somatic tissues compared to UASt constructs (DeLuca and Spradling, 2018). We found that by using UASp-2xMCP2xeGFP, expression was lower in the somatic tissues we tested including the wing disc, brain and ovarian follicle cells. As this avoided accumulation of the strongly fluorescent MCPeGFP aggregates, we used these transgenes in our further experiments.

We used a fly line with *ptc-*Gal4 driving UASp 2xMCP2xeGFP to visualise MS2 transcription in larval and adult tissues. *ptc-*Gal4 is expressed in the wing disc, follicle cells and brain (Weaver et al., 2020). We focussed on the *Daughters against decapentaplegic* (*Dad*) gene which is expressed in the ovary and brain and studied *Dad* transcription in these tissues by using a MiMIC insertion to create a fly line with 128xMS2 loops in the first coding intron (Fig. 5A). *ptc*-Gal4; UASp-2xMCP2xeGFP females were crossed to *Dad*-128xMS2 males. In dissected tissue we live imaged the follicle cells (Fig. 5B) and detected transcription of *Dad*-128xMS2 in the anterior follicle cells of stage eight egg chambers (Fig. 5C, Movie 6). This is consistent with previous expression data using a *Dad-lacZ* reporter line (Shravage et al., 2007). Moreover, visualisation of nascent transcription sites in these tissues using smFISH with endogenous *Dad* probes and MS2 probes confirmed that MS2 detection accurately depicts the endogenous transcription of this gene (Fig. S3A).

**Figure 5.**
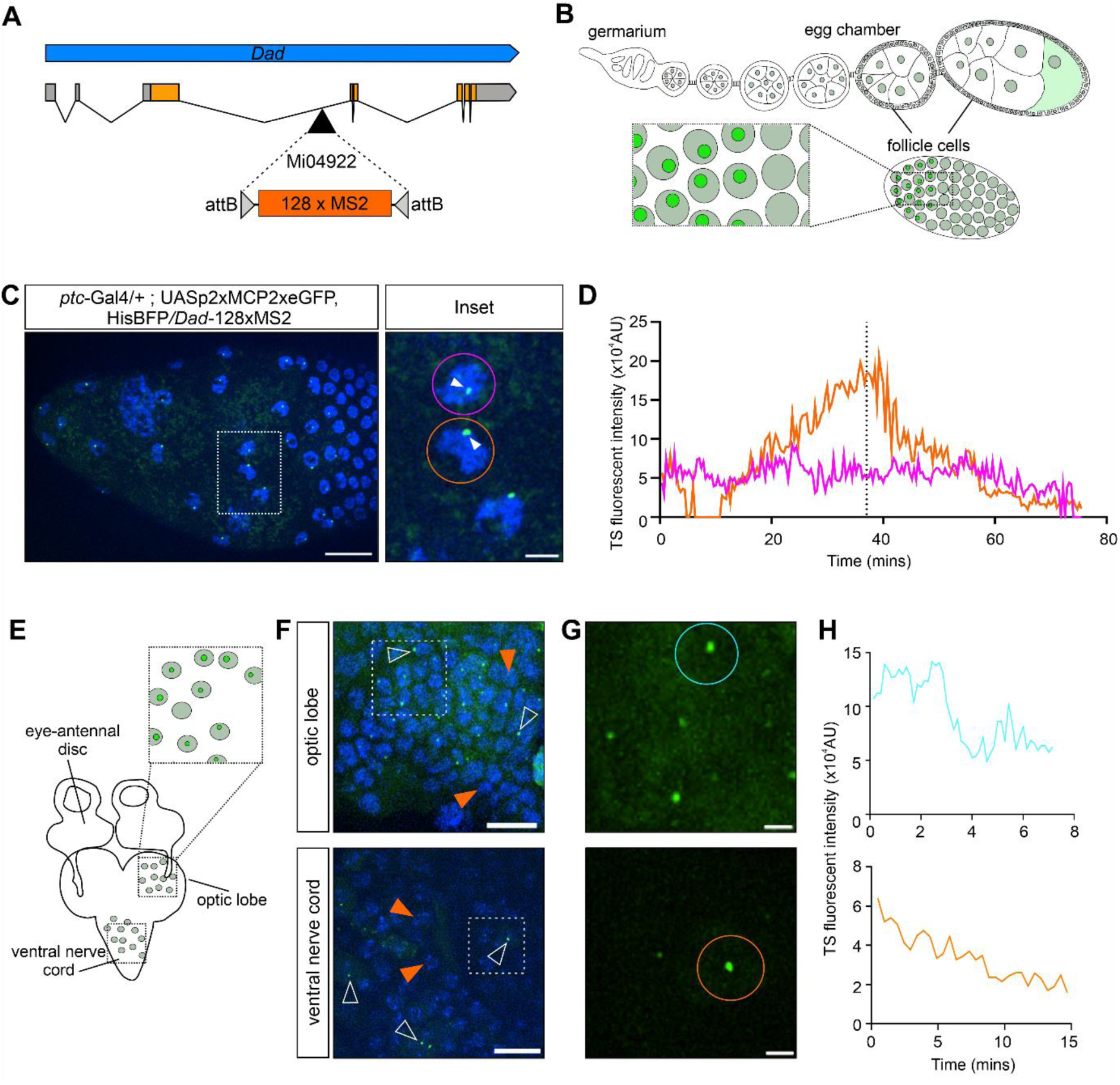
Visualising nascent gene expression in the ovary and larval brain using the Gal4-UAS system. (A) Schematic of the 128xMS2 loop array inserted into the Mi04922 insertion in the first coding intron of the *Dad* gene. (B) Schematic showing the female ovary and the follicle cells that surround the egg chambers and oocyte. (C) Still from a live imaging movie of a stage eight egg chamber from a *ptc-Gal4/+; UASp-2xMCP2xeGFP/Dad-128xMS2* female, anterior is to the left. *Dad*-128xMS2 TSs are detected in the anterior follicle cells with a region of expressing cells marked by the white box. Scale bar is 20 µm. Inset: A higher magnification image of the follicle cells highlighting two cells (purple and orange circles) with transcription sites (arrowheads). Scale bar is 5 µm. (D) Fluorescent intensity traces from the two cells highlighted in the inset. The colours denote the different cells in the inset in (C). (E) Schematic showing the third instar larval brain and eye-antennal discs highlighting the regions of the optic lobe and ventral nerve cord that are imaged live. (F) Stills from live imaging movies of cells from *ptc*-Gal4> UASp-2xMCP2xeGFP and *Dad*-128xMS2 larval brains. A number of cells have active transcription sites (white arrowheads), and examples of non-transcribing cells are also present (orange arrowheads). Scale bar is 10 µm. (G) Higher magnification of the outlined regions in (F). Scale bar is 2µm. (H) Transcription traces from the circled brain cells in (G).

Live imaging over a period of an hour showed that transcription was maintained in the tissue throughout the imaging period. Transcriptional traces of two cells showed that they had different active transcription times and off periods (Fig. 5D). The magenta cell maintained a stable level of transcription throughout the imaging window whereas the orange cell had a period where transcription was off for ∼5 minutes and then had a higher level of transcription between 20 and 45 minutes compared to the magenta cell. These data demonstrate that transcription can be imaged in ovarian follicle cells over a significant time and suggest that there is heterogeneity in the transcriptional activity between cells.

We next live imaged dissected third instar larval brains (Fig. 5E) and observed *Dad*-128xMS2 TSs in cells of the optic lobe and ventral nerve cord (Fig. 5F, Movie 7). Expression of *Dad* in the larval brain is consistent with previous reports of a *Dad-lacZ* reporter expressed in the ventral nerve cord (Vuilleumier et al., 2019). Tracking of TSs within these cells showed dynamic transcription across the short 15-minute imaging period (Fig. 5H). Together, these data show the feasibility of studying transcription dynamics over varying time periods of minutes up to hours in living tissues of the ovary and larvae.

In addition to the UASp lines we also tested an existing stock with the *Hsp83* promoter driving MCPeGFP with a nuclear localisation signal (Forrest and Gavis, 2003) to visualise MS2 tagged mRNAs in larval and adult tissues. *Dad-*128xMS2 males were crossed to Hsp83MCPeGFP females to detect transcription of *Dad*-128xMS2 in the cells of the ovary (Fig. S3) and wing disc (Fig. S4). We also detected *Dad*-128xMS2 TSs in the anterior follicle cells of the ovary as observed with *ptc-*Gal4*>*UASp-2xMCP2xeGFP and tracked these nuclei for more than 2 hours (Fig. S3B-C).

Live imaging of *Dad-*128xMS2 in the wing disc focussed on the squamous cells on the surface of the wing disc pouch (Fig. S4A). TSs, with stronger signal than the low level of free MCPeGFP in nuclei, were observed in many cells (Fig. S4B) and this expression was confirmed in fixed tissue using smFISH (Fig. S4D). In addition, we observed nonspecific accumulation of MCPeGFP in a larger structure within the nucleus (Fig. S4B), which appears to be the nucleolus based on colocalization of the GFP signal with staining for the nucleolar marker fibrillarin in fixed wing disc tissue (Fig. S4E). Therefore, it will be important to include a control without MS2 when using the Hsp83MCPeGFP stock, to confirm the transcription site is being labelled in addition to the nucleolus (Fig. S4D). We rarely observed the nascent transcription site localised near to the nucleolar region and were able to track the TSs in individual nuclei over time, which showed bursts of transcription (Fig. S4C). Overall, these data show that the MiMIC-based approach can be used to insert MS2/PP7 loops into endogenous genes, facilitating the visualisation of transcription live for long periods of time in different *Drosophila* tissues.

## Discussion

Here we have developed a new set of *Drosophila* stocks that allow MS2/PP7 stem loops to be inserted into protein coding genes and ncRNAs to visualise nascent transcription *in vivo*. We have exploited a previously reported MiMIC-based gene tagging strategy that relies on RMCE (Nagarkar-Jaiswal et al., 2015a) and combined it with our new stocks so that insertion of the stem-loops relies only on a simple crossing scheme and PCR screening. As proof-of-principle, we have used different MiMIC stocks to show how this approach can be used to visualise transcription of different genes and a lncRNA in distinct tissues including the embryo, larval wing disc, brain and ovary. The versatile options for visualising transcription presented here, e.g. the new coat protein lines, and the ease of use of this MiMIC based approach will benefit many researchers in the *Drosophila* research community.

Current methods to insert MS2 or PP7 loops into endogenous genes using CRISPR can be time-consuming and difficult. CRISPR requires many cloning steps or the design plus synthesis of gene strings, followed by embryo microinjections, screening for successful CRISPR events, verification of the insertion and establishment of the stocks. ∼10 weeks is needed for stem-loops to be inserted into a gene locus (Gratz et al., 2015; Hoppe and Ashe, 2021a; Yu et al., 2021). While transgenic lines can be more rapidly generated, these may not contain all the regulatory sequences required to fully recapitulate the endogenous transcription dynamics. The method presented here can routinely be performed in ∼5 weeks as the process simply involves crossing fly stocks followed by verification of the insertion by PCR, while avoiding the more costly and time-consuming steps of cloning and microinjection.

Combining these new reagents with the MiMIC insertion collection opens up the possibility for at least 4367 genes to be targeted throughout the genome. One obvious limitation to the system presented here is that a MiMIC insertion is required. If the specific gene of interest does not contain a MiMIC insertion, then a CRiMIC insertion could be generated. CRiMIC insertions are MiMIC insertions inserted into the genome via CRISPR mutagenesis so that they are targeted to genes rather than being randomly inserted (Lee et al., 2018). Generating a CRiMIC has the advantage that it would also allow fluorescent protein tagging in addition to insertion of stem-loops (Lee et al., 2018). Many CRiMIC insertions are available from the Bloomington stock centre, with this number likely to rise in the future.

One benefit of utilising the MiMIC-based approach described here is that there is an abundance of target sites located in the UTRs and introns of genes, which are the preferred sites of MS2/PP7 stem loop insertions to avoid disrupting coding sequences. MS2/PP7 stem loops inserted in the 5’ UTR, 3’ UTR or an intron have all been used to study transcription dynamics previously (Forbes Beadle et al., 2023; Garcia et al., 2013; Lucas et al., 2013; Whitney et al., 2022). Each position has advantages and disadvantages relating to the study of promoter states (Ferraro et al., 2016). While insertion into the 5’ UTR results in a high signal to noise, the persistence of fluorescence hampers the direct detection of transient off states. Conversely, with stem loops in the 3’ UTR the fluctuations in signal better represent the promoter states but the signal is much weaker and more sensitive to background coat protein levels. If the MS2/PP7 loops are inserted into the intron, then the fluorescent signal depends on both the promoter state and the efficiency of splicing (Ferraro et al., 2016). However, data on intron half-lives in *Drosophila* S2 cells are available, which show that the median intron half-life is only 2 minutes (Pai et al., 2017).

Our data demonstrate how insertion of PP7 and MS2 sequences into the same MiMIC site can be used to simultaneously visualise both *pxb* alleles in the same embryo. This will, for example, allow cell to cell variation in transcription to be assigned to intrinsic versus extrinsic variation (Falo-Sanjuan et al., 2019). In addition to studying transcription dynamics, MS2/PP7 tagging of genes and detection of transcription sites can be used to mark cell populations and development events. For example, an *engrailed-MS2* reporter has been used to identify and study parasegment boundaries during germ band extension in living wildtype and mutant embryos (Sharrock et al., 2022). Finally, the 128xMS2 cassette allows single mRNAs to be visualised in the cytoplasm (Dufourt et al., 2021; Vinter et al., 2021), facilitating the study of mRNA localisation.

MiMIC-based MS2/PP7 insertion can also be used to study the transcription and localisation of ncRNAs. Flybase reports ∼200 MiMIC-containing lncRNA stocks available across the genome. Previous studies inserted 24xMS2 loops into the *iab-8* ncRNA from the bithorax complex (Arib et al., 2015) and 6xMS2 into the *roX1* ncRNA (Apte et al., 2014). However, these stocks were only used to visualise the transcription sites in fixed tissues. Here, by inserting 24x PP7 sites into a MiMIC in *roX1*, we have visualised *roX1* transcription in embryos during nc14. Our data show that *roX1* transcriptional activation in individual cells is very synchronous in the embryo, and that it is only transcribed transiently. How *roX1* transcription kinetics relate to its function in dosage compensation at different times and in distinct tissues is an interesting question that can be investigated with this stock in the future. Allele-sensitive single-cell RNA sequencing has revealed that mammalian lncRNAs have a high cell to cell variability and lower abundance due to a reduced burst frequency compared to mRNAs in the same tissue (Johnsson et al., 2022). It will also be interesting to determine how transcriptional burst parameters from ncRNAs compare to those from mRNAs during *Drosophila* development. The new coat protein lines generated here will facilitate studies of transcription of a protein coding gene and lncRNA in the same cells during *Drosophila* development.

The vast majority of studies of live transcription in *Drosophila* have been performed in the early embryo. In this study, we have shown how MS2 imaging can be used to visualise and quantitate live transcription of the *Dad* gene in the ovary, larval brain and wing disc. In the ovary, long-term imaging of *Dad-*128xMS2 TSs showed heterogeneity in the transcriptional responses in individual cells, whereas bursty *Dad* transcription was detected in wing disc cells. To express the coat proteins for live imaging in the ovary and larval brain we found that the use of GAL4 drivers with the UASp expression system, which is weak in somatic tissues (DeLuca and Spradling, 2018), allowed the best signal to noise detection of transcription sites. The *Hsp83* promoter is another useful option, although this resulted in MCPeGFP accumulation in the nucleolus in some cells. In contrast, we found that the very high levels of fluorescent coat proteins obtained with the GAL4/UASt system caused accumulation in cells that impeded detection of live, nascent TS signals. Keeping tissues alive and transcriptionally active during live imaging is another consideration. However, many protocols already exist that allow *ex vivo* imaging of tissues such as we have used here in the larval discs (Dye, 2022) and the ovary (Wilcockson and Ashe, 2021). As these tissues have longer developmental time windows than the early embryo this will allow new features of promoter dynamics and transcription to be studied. For example, it is possible that multi-scale bursting will be observed in which the promoter fluctuates on both short (minutes) and long (hours) time scales, as has been described in human cells (Tantale et al., 2016).

In summary, the MiMIC-based approach for inserting MS2/PP7 stem loops and new coat protein lines that we have described here will be useful to many researchers interested in probing transcriptional dynamics and/or marking specific cell types in their tissue of interest. Moreover, such studies of *in vivo* transcriptional dynamics can be combined with other techniques such as single cell transcriptomics to provide wholistic and time resolved models of transcription in varying developmental contexts and in response to different cues.

## Materials and Methods

### Fly stocks

All stocks were routinely maintained at 18°C and experiments performed at 25°C unless otherwise specified on standard fly food media (yeast 50g/L, glucose 78g/L, maize 72g/L, agar 8g/L, 10% nipagen in EtOH 27mL/L and propionic acid 3mL/L). All stocks used and produced in this manuscript are listed in Table 1.

**Table 1.**
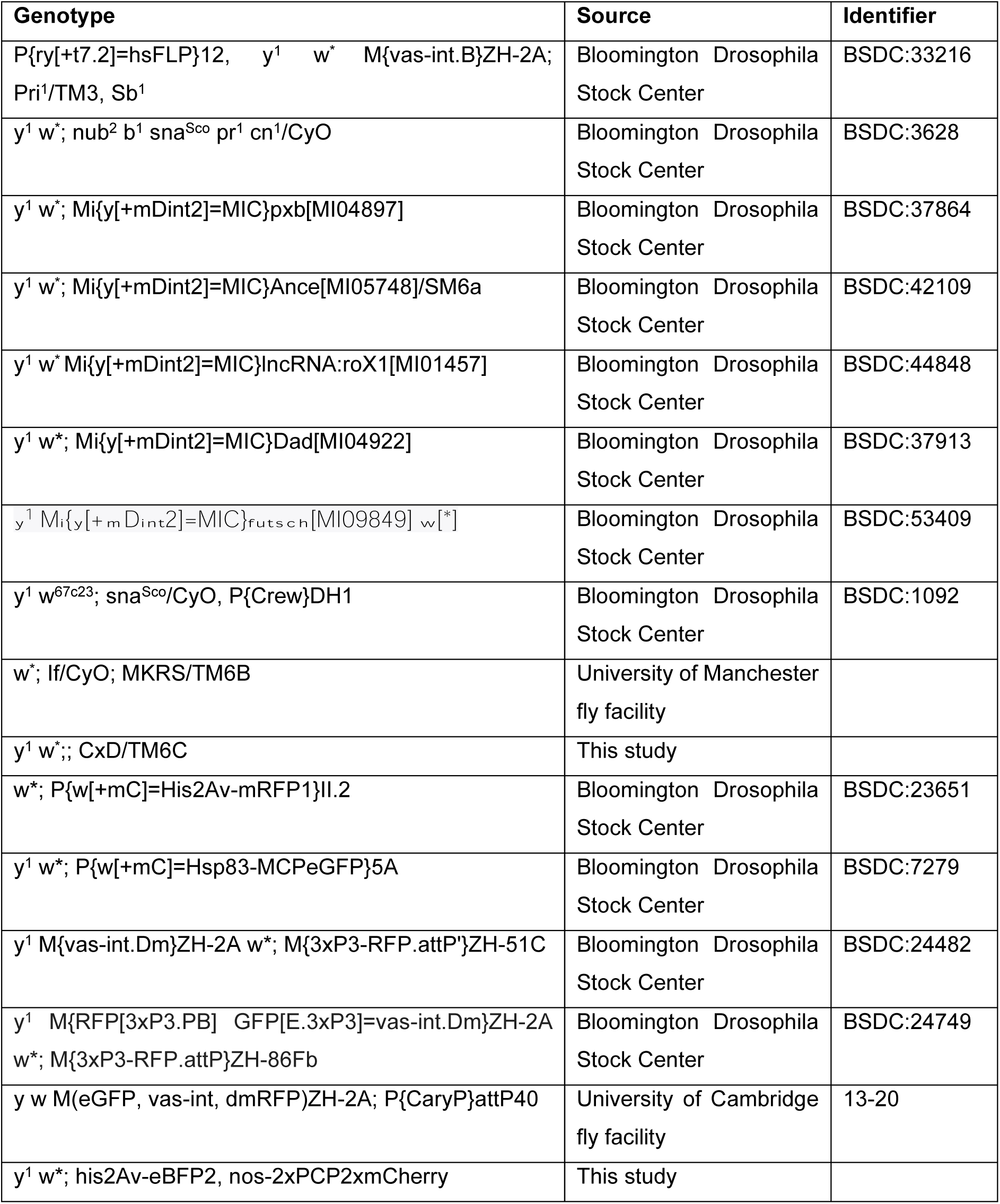
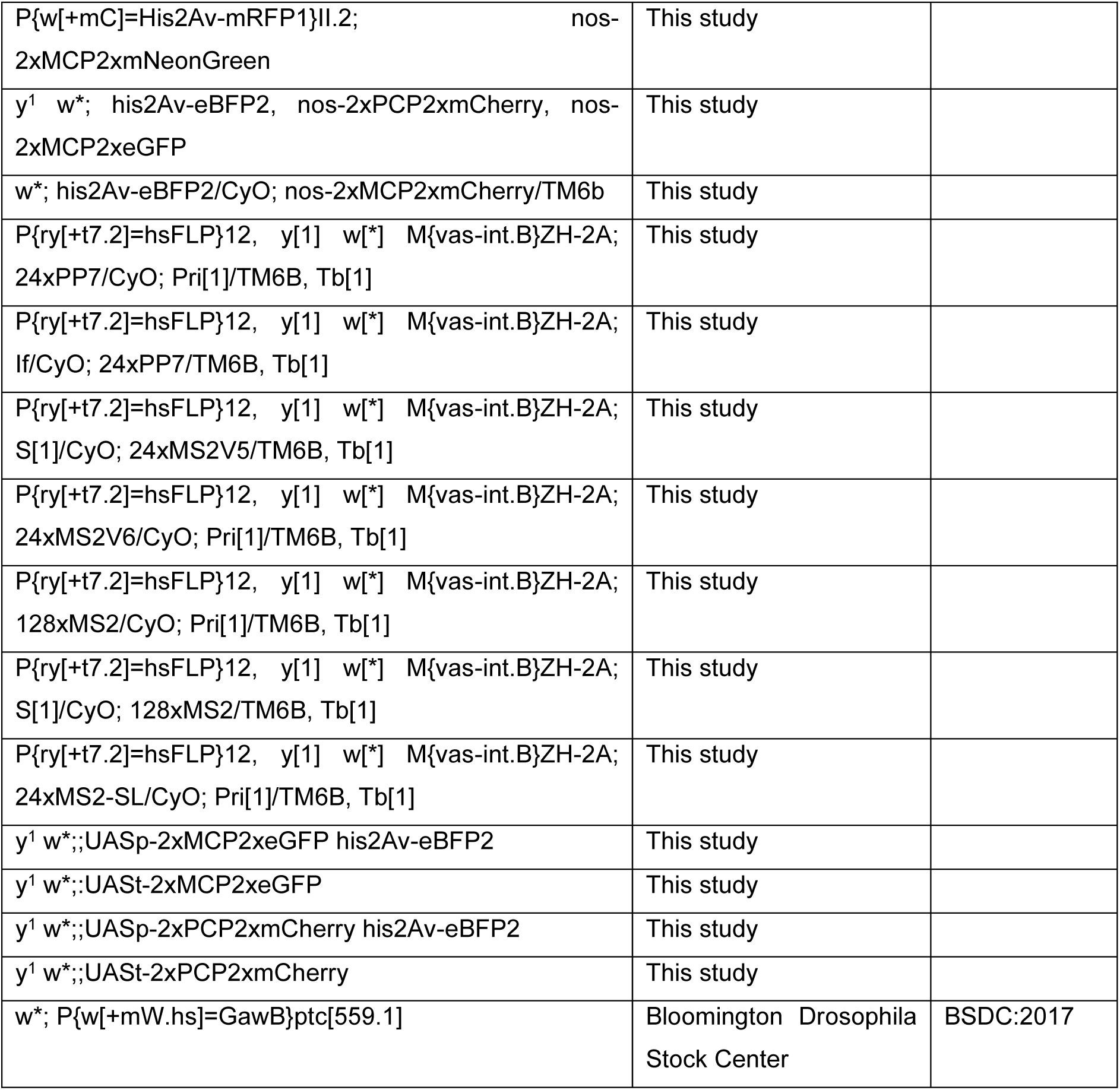
Fly stocks used and made in this manuscript.

### Cloning

#### pw35 plus attB and loop donor vectors

To construct the loop donor cassette vectors the pBS-KS-attB1-2 cloning vector (JN222909.1, Addgene #61255) was cut with NheI and NsiI to obtain a 264bp fragment with the inverted attB sites, and this fragment was inserted into pw35 (DGRC Stock 1168) that had been linearised with PstI and AvrII to form a pw35-attB1-2 vector. Each of the repetitive loop sequences were cut out of their original vectors and inserted into the pw35-attB1-2 vector. Vectors pCR4-24xMS2SL-stable (Addgene #31865), pBlueScript-24xPP7 (Fukaya et al., 2016) and pET264-pUC 24xMS2V6 Loxp KANr Loxp (Addgene #104393) were digested using BamHI and BglII to obtain 24xMS2-SL, 24xPP7 and 24xMS2V6 respectively. These fragments were inserted into the pw35-attB1-2 vector which had been digested with BglII. For 24xMS2V5 the cassette was PCR amplified out of the original vector (Addgene #84561) using primers containing additional PstI flanking sequences, digested with Pst1 and inserted into pw35-attB1-2 cut with PstI. For 128xMS2 cloning, an extra MCS containing an NheI site was first cloned into pw35-attB1-2 using PstI and BamHI The 128xMS2 loops from pMK123-128MS2(XbaI) (Tantale et al., 2016) were excised using XbaI and BamHI and inserted into the BamHI and NheI sites of pw35 plus attB.

#### pCasper-attB-nos-2xMCP2xeGFP, pCasper-attB-nos-2xMCP2xmCherry and pCasper-attB-nos-2xPCP-2xmCherry

A vector modified from pCasper (DGRC 1213) with an added attB site was used to insert the 948bp nanos promoter region (3R:19156372-19157319) into the EcoRI site and the 1295bp tubulin 3’UTR and downstream genomic region (3R:7088576-7089870) in the XbaI site. This pCasper-attB-nos-tubulin 3’UTR-attB vector (Vinter et al., 2021) was used to clone 2xMCP and 2xPCP with 2xmCherry and 2xeGFP. 2xMCP was derived from pHsp83-NLS-HA-2xMCP2xTagRFP-T (Addgene #71242) (Halstead et al., 2015) and cloned 5’ to 2xeGFP or 2xmCherry amplified from pHsp83-NLS-HA-2xPCP-2xGFP vector (Addgene #71243) (Halstead et al., 2015) or pTV Cherry (DGRC Stock 1338) into KpnI/BamHI of pCasper-nos-tubulin 3’UTR-attB, with a linker between the two GFP fluorophores using Infusion (Clontech, Takara Biosciences). 2xPCP from the pHsp83-NLS-HA-2xPCP-2xGFP vector (Addgene #71243) (Halstead et al., 2015) was cloned 5’ to 2xmCherry from pTV Cherry (DGRC Stock 1338) into KpnI/BamHI of pCasper-nos-tubulin 3’UTR-attB, with a G_4_SG_4_S_2_RM linker between the two fluorophores using Infusion (Clontech, Takara Biosciences).

#### pCasper-attB-nos-2xMCP2xmNeonGreen

The NLS-2xMCP from pHsp83-NLS-HA-2xMCP2xTagRFP-T (Addgene #71242) (Halstead et al., 2015) was cloned into KpnI/SpeI of pCasper-attB-nos-tubulin 3’UTR. The NLS was removed from the 2xMCP by digesting with KpnI and NheI and repaired using primer annealing. 2xmNeonGreen sequences were added from pCasper_nosP_scFVmNeonGreen-GB1-NLS_tub3UTR (Vinter et al., 2021) and inserted into the SpeI site downstream of the 2xMCP using the primers in Table S1.

#### pCasper-attB-nos-2xPCP2xmCherry-his2Av-eBFP

The his2Av-eBFP insert from nanos-SV40NLS-mCherry-PCP-his2Av-eBFP (Fukaya et al., 2017) was subcloned into the SacII site of pCasper-nos-2xPCP-2xmCherry-tubulin 3’UTR.

#### pUASp-attB-2xMCP2xeGFP-his2Av-eBFP, pUASp-attB-2xPCP-2xmCherry-his2AveBFP, pUASt-attB-2xMCP2xeGFP and pUASt-attB-2xPCP-2xmCherry

The 2xPCP-2xmCherry-tubulin 3’UTR and 2xMCP2xeGFP-tubulin 3’UTR were subcloned from the above pCasper-attB vectors into KpnI/NdeI of UASp-attB and EcoRI/NotI of pUASt-attB. The his2AveBFP insert was subcloned into SacII of pUASp-attB-2xPCP2xmCherry and pUASp-attB-2xMCP2xeGFP.

### Microinjections and transformant screening

Plasmid DNA was prepared for microinjection using a Qiagen Plasmid Mini or Maxiprep kit (Cat #12123 or #12163) according to the manufacturer’s instructions and including an extra PB wash step. DNA was diluted to the required injection concentrations in nuclease free water and all injections were performed by the University of Manchester or Cambridge fly facilities. For the loop donor cassettes (24xMS2-SL, 24xPP7, 24xMS2V5, 24xMS2V6 and 128xMS2) and pCasper-nos-2xMCP2xmCherry and pCasper-nos-2xPCP2xmCherry the plasmid DNA was injected into *w*^1118^ *Drosophila* embryos with P element helper plasmid (DGRC Stock 1001) at 0.8 µg/µl and 0.5 µg/µl respectively. Successful random P element insertions were selected by screening for red eyed progeny (*w*^+^) after crossing the injected individuals to *w*^1118^. Transformants were crossed to a second and third chromosome balancer stock (w*; IF/CyO; MKRS/TM6B) to map them to chromosomes and subsequently the P element was mapped using inverse PCR (Bellen et al., 2004). At least one insertion on the second and third chromosome was kept for subsequent crossing. Transformants were then crossed into the P{hsFLP}12, *y*^1^ *w*\* M{vas-int.B}ZH-2A; S^1^/CyO; Pri^1^/TM6B, Tb^1^ (BSDC:33216) background and maintained at 18°C to avoid leaky expression of the hsFLP that can occasionally remove the loop donor cassette. Escaper flies with *white*^-^ eyes which occasionally flip out the loop cassette flanked by FRT sites were routinely removed from stocks to maintain the loop cassette.

pCasper-attB-nos-2xPCP2xmCherry-his2AveBFP was inserted into attP40 (stock 13-20 from the University of Cambridge fly facility: *y w M(eGFP, vas-int, dmRFP)ZH-2A; P{CaryP}attP40*), and pCasper-attB-nos-2xMCP2xeGFP was inserted into attP51C (BDSC:24482: *y^1^ M{vas-int.Dm}ZH-2A w*; M{3xP3-RFP.attP’}ZH-51C)*. These were recombined to generate a line expressing both coat proteins and histone marker on the 2^nd^ chromosome his2av-eBFP, 2xPCP2xmCherry, 2xMCP2xeGFP.

pUASp-attB-2xMCP2xeGFP-his2AveBFP, pUASp-attB-2xPCP-2xmCherry-his2AveBFP, pUASt-attB-2xMCP2xeGFP and pUASt-attB-2xPCP2xmCherry were inserted into attP86Fb (BSDC: 24749) on the 3rd chromosome. The X chromosome 3xP3-RFP and -GFP markers were selected against after selecting for successful transformants and the landing site 3xP3-RFP was removed using cre recombinase either before or after injections.

2xMCP2xmNeonGreen was randomly inserted using P element mediated transgenesis as above into the his2AvRFP line (BSDC:23651) and a single third chromosome insertion stock was found after screening using inverse PCR P element mapping.

See Table 1 for the stock combinations that were made from these fluorescently tagged coat protein insertions. All stocks made in this study are available upon request.

### Genomic DNA extractions and PCR screening

Genomic DNA was extracted from 10-15 adult flies. Screening for RMCE events was performed via PCR as in (Venken et al., 2011) using MIL_F and MIL_R primers in combination with attB_MS2_2 and a primer specific for the loop type. Primers were designed to overlap the attB/loop sequence boundary to avoid the most repetitive sequences of the intervening stem loop sequences. All primer sequences are listed in Table S1. PCR cycling conditions were as follows 95°C for 2 min, then 30 cycles of 95°C for 30 secs, annealing at 60°C for 30 sec, 30 sec extension at 72°C using GoTaq polymerase (Promega # M7122). A final extension of 72°C completed the PCR reactions. As the loops and the attL/R scars are composed of repetitive sequence, PCR based methods of screening using primers in these regions can potentially lead to laddering and nonspecific products whereby the newly synthesised PCR products can produce a range of sizes by acting as primers in the subsequent rounds of primer annealing and amplification. We recommend using a PCR cycling protocol with a short annealing time and a low number of cycles as described here in the first round of screening.

### Crossing scheme and heat-shock

For second and third chromosome targeting, 30 virgin adult females containing the loop cassette were crossed to 15 males with the MiMIC insertion in the gene of interest in single vials. For X chromosome insertions, 30 virgin adult females containing the MiMIC insertion were crossed to 15 males with the loop cassette in single vials. After allowing flies to lay on the food for 3 days (Day 1-3) the flies were transferred onto fresh food. Following removal of adult flies on day 3 the vial containing embryos and larvae aged 1-3 days old was immediately heat shocked in a water bath at 37°C for 30 minutes. The same vial was then heat-shocked again on day 5 and 7. All crosses were kept at 18°C until heat-shock and the vials with progeny were shifted to 25°C after heat shock and until adults emerged. A single 30-minute heat shock on day 3 was also successful.

Once adults have emerged the mosaic eyed *y*+ progeny are then crossed to a balancer stock. In the next generation, *yw* males should be selected, balanced and screened for direction of the stem loop insert for autosomal insertions. For insertions on the X chromosome, *yw* males must be selected for and additionally *ry*+ should be selected against. The vasa phiC31 integrase insertion site is also marked by a 3xP3-RFP which can be selected against instead of *ry+* in the progeny at the final step if preferred.

For X chromosome targeting the donor cassette flies which have the heat shock-FLP recombinase and vasa-integrase must be crossed in from the male parent as the female parent must provide the MiMIC insertion. Therefore, in the case of X chromosome targeting recombination can occur between the MiMIC containing *yellow*+ locus and the yellow-mutation at the endogenous locus in the parental loop donor cassette stock. This may result in *yellow*-progeny emerging at the end of the crossing scheme that do not have the loops inserted into the MiMIC and are instead products of recombination between the two loci. In our hands we found that setting up many crosses allowed us to obtain enough progeny to get positive hits via PCR screening even when accounting for recombination between the *lncroX1* or *futsch* locus and *yellow*. More crosses will need to be set up the further away the targeted X chromosome gene is from the *yellow* gene. In some rare cases it may be difficult to target the gene of interest if it sits far from the *yellow* locus on the X chromosome, but we anticipate that with enough progeny a successful insertion will occur.

### Embryo and tissue smFISH and immunofluorescence

Embryos were laid on apple juice agar plates supplemented with yeast paste in a 25°C incubator for 2 hours then aged for another 2 hours. Embryos were then dechorionated in 50% bleach solution (2.5% final concentration of sodium hypochlorite solution diluted in water) and washed thoroughly in ddH_2_O. Embryos were transferred into 3ml of fixation buffer (1.3xPBS, 67mM EGTA pH8) in a scintillation vial and 1ml of 37% formaldehyde and 4ml of heptane was added. Embryos were fixed for 20 minutes with shaking at 300rpm as described (Kosman et al., 2004). After settling, the lower phase was removed and 8ml methanol was added and vortexed for 1 minute. The upper phase was then removed, and embryos were washed in methanol 3 times before storage at -20°C.

Dissected tissues were fixed in 4% formaldehyde for 20 minutes on a rotating wheel. After fixation, three 5-minute washes each in 25%, 50% and 75% methanol in PBT (1x phosphate buffered saline with 0.05% Tween20) were performed. Tissues were washed in 100% methanol for 10 minutes and then transitioned back to 25% methanol in PBT using three 5-minute washes. Tissues were processed immediately following fixation for smFISH.

smFISH and immunostaining was performed on embryos as described in (Vinter et al., 2021) with the following modifications. Ovary tissues were incubated overnight at 4°C with primary antibodies: rabbit anti-fibrillarin (Abcam #ab5821) and goat anti-GFP antibody (Abcam #ab6673). Secondary antibody incubation was for 2 hours using Alexa Fluor antibodies: donkey anti rabbit 647 (Thermo Fisher Scientific #A-31573) and donkey anti goat 488 (Thermo Fisher Scientific #A-11055). Just before mounting, ovaries were pipetted up and down with a P1000 tip to separate the ovarioles before mounting on slides in Prolong Diamond Antifade Mountant (Thermo Fisher Scientific #P36961). smFISH probe sequences are listed in Table S2.

### Live imaging Microscopy

Embryos were laid on apple juice agar plates supplemented with yeast paste in a 25°C incubator for approximately 1 hour and aged if required to just before the desired age. Embryos were collected and dechorionated in 50% bleach solution (2.5% final concentration of sodium hypochlorite solution diluted in water). FluoroDish tissue culture dishes with a cover glass bottom (World Precision Instruments #FD3510-100) were coated with a thin layer of heptane glue and embryos were mounted onto the heptane glue coated dishes. A drop of Halocarbon oil (7:1 mix of 700:27 Halocarbon oil; Sigma #H8898 and #H8773) was applied over the embryos to keep them from drying out during the imaging period.

For live-imaging of wing discs, wandering third instar larvae were dissected in Grace’s insect media (Sigma Aldrich #G8142-500ML) supplemented with 5% FBS 1% Pen/Strep according to (Dye, 2022). Dissected tissues were then transferred into FluoroDish tissue culture dishes in Grace’s media for imaging. Ovaries were dissected from 5-7 day old female flies and mounted onto FluoroDish tissue culture dishes as described (Wilcockson and Ashe, 2021).

Images were collected on an Andor Dragonfly200 spinning disk inverted confocal microscope with a Piezo stage using either a 40x/1.30 Super fluor or 100Xx/1.4 Plan Apo VC objective. Samples were excited using a combination of 405nm (5-10%), 488nm (10%) and 561nm (10-20%) diode lasers and 450nm DAPI, 525nm GFP or 600nm RFP filters respectively. The laser power and exposure times were set as to not overexpose or bleach the fluorescent signal and differed between genotypes. Each channel was collected sequentially with the iXon EMCCD (1024 X 1024) camera with a 100-150ms exposure time per channel. For each movie a total of 30-50 Z stacks at 0.5µm spacing were collected continuously using the fastest setting yielding a total Z size of 15-25µm at a time resolution of between 20-25 seconds on average.

### Fixed imaging Microscopy

Confocal Z stack images of fixed embryos and wing disc tissues were collected using an Andor Dragonfly200 spinning disk inverted confocal microscope as above with 2 or 4x averaging and a combination of 405nm (5-10%), 488nm (10%), 561nm (10-20%) and 637 (15%) diode lasers and 405nm DAPI, 488nm GFP, 561nm RFP or 637 Cy5 filters respectively using system optimised step-size. Confocal Z stack images of ovaries were collected using a Leica gSTED SP8 inverted confocal microscope using a HC PL APO CS2 100x/1.40 oil objective with pinhole of 1 airy units, 3.5x zoom, bidirectional scan speed 400Hz, with an 8bit image of size 2048 X 2048 pixel and 4x line averaging at a system-optimised step size. Images were collected by illumination with a white light laser at 70% with the following detection settings: PMT DAPI excitation at 15% 405 nm (collection: 415 to 482 nm); Hybrid Detectors: Quasar 570 excitation at 20% 548 nm (collection: 565 to 631 nm), Quasar 670 excitation at 20% 647 nm (collection: 657 to 735 nm) with 1 to 6ns gating.

### Image processing and analysis

Fiji (Image J) was used for image processing to crop and make maximum projections of confocal images and timelapse movies. GraphPad Prism 9 and R studio software was used for plotting and data analysis.

### Transcription site detection and tracking

Imaris 10.1(Bitplane) was used to segment nuclei and transcription sites using the “surface” function for nuclei and “spot” function for TS. Spots had an XY diameter of 1 and Z diameter of 2. Autoregression tracking of the nuclei used a maximum distance of 5µm and maximum gap size of 3. Background spots of the same size as the TS spots were added at every third timepoint. Nuclei and spot statistics were run through the sass algorithm (Hoppe and Ashe, 2021b) that assigns spots to nuclei over time with background subtraction to give background corrected fluorescent intensity of each TS over time assigned to its nearest nuclei.

For movies of follicle cells and wing discs the TS were detected as “spots” as described above but with tracking using autoregressive motion and a maximum distance of 1.5µm and gap size of 3. Background intensity was calculated by doubling the volume of each of the spots over time and subtracting the spot intensity sum from the double spot volume intensity sum. This background intensity was then subtracted from each spot intensity sum for each transcription site “spot” to give a background subtracted spot sum intensity at each time point.

## Acknowledgements

We thank Nathan Garnham for making pCasper-nos-2xMCPneonGreen, Hadi Boukhatmi for suggesting using the Hsp83MCPeGFP flies, Sanjai Patel at the University of Manchester Fly Facility and Cambridge University for generating the transgenic flies, and the University of Manchester Bioimaging Facility for support.

## Author contributions

Conceptualisation: L.F.B., H.L.A.; Investigation: L.F.B., C.S.; Writing - original draft: L.F.B., H.L.A.; Writing - review & editing: L.F.B., C.S., H.L.A.; Supervision: H.L.A., Funding acquisition: H.L.A.

## Data Availability

All relevant data can be found within the article and its supplementary information.

---

**Figure S1.**
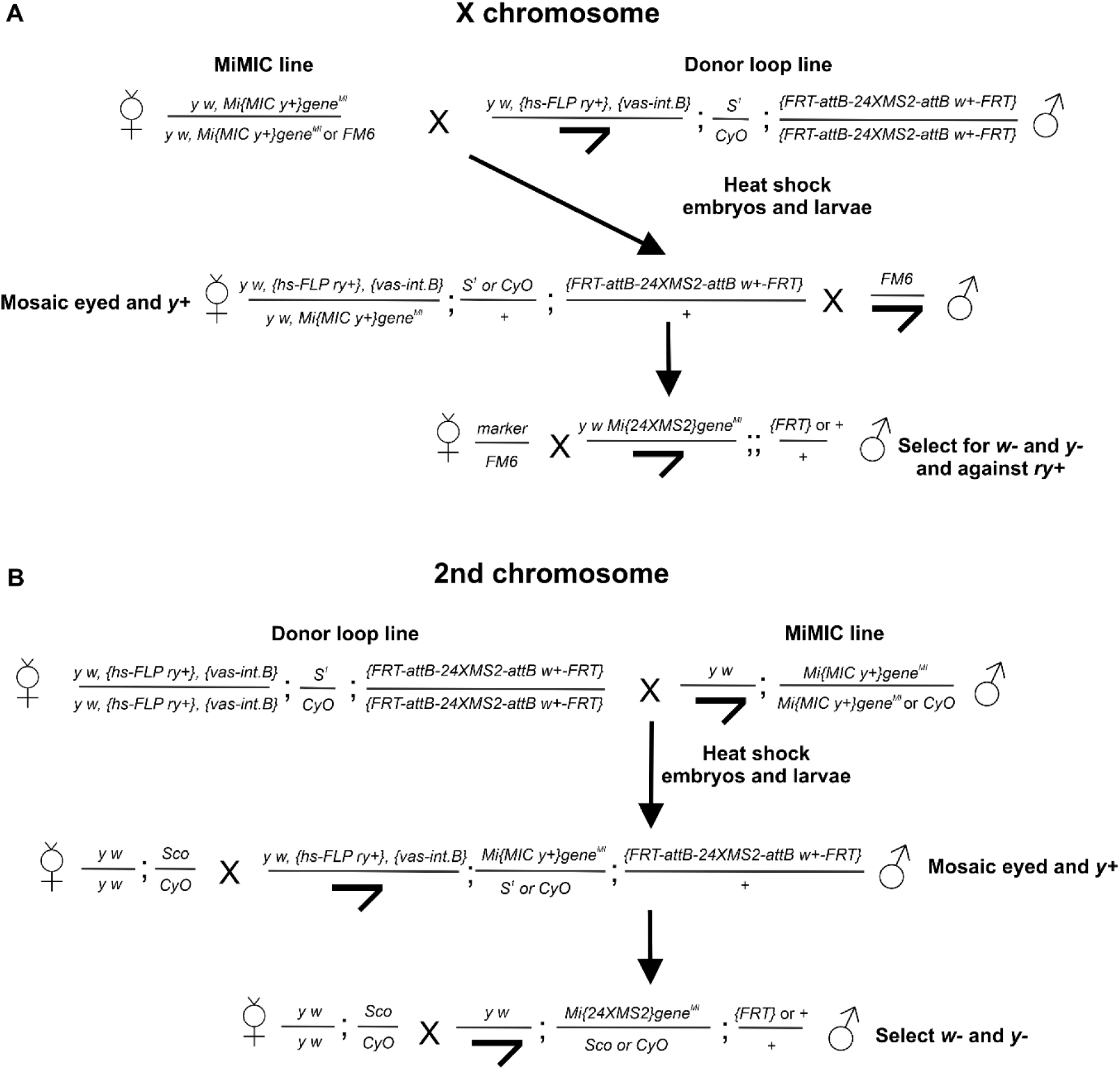
Crossing scheme for inserting loop sequences into genes on the X and second chromosome. (A, B) Crossing scheme for insertion of 24xMS2 loops into a MiMIC on the X (A) and second (B) chromosome.

**Figure S2.**
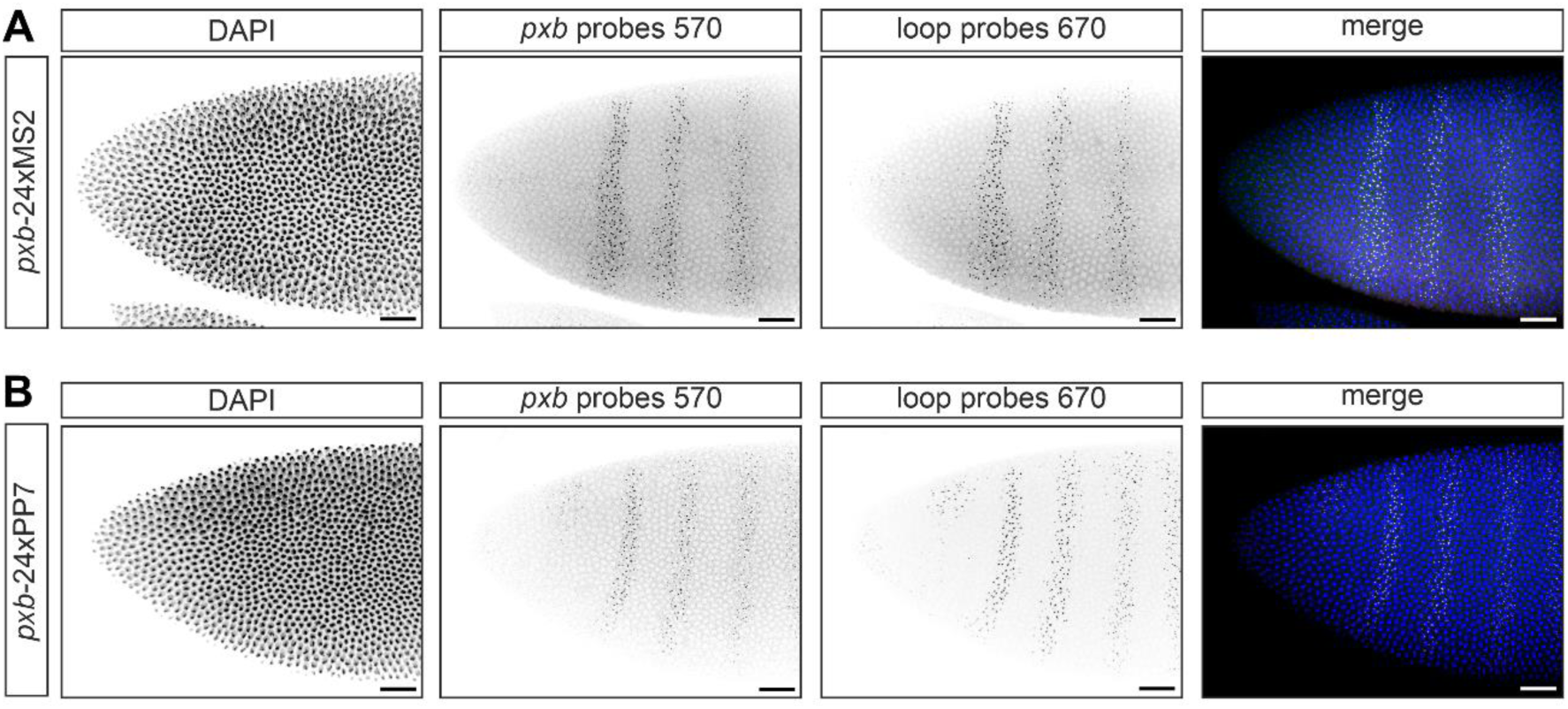
*pxb* transcription in embryos endogenously tagged with 24xMS2 or 24xPP7 loops. (A) Confocal images of *pxb*-24xMS2 embryos showing the anterior stripes of the *pxb* expression domain visualised by smFISH staining with *pxb* probes (570 channel, green) and MS2 loop probes (670 channel, magenta). Nuclei are labelled with DAPI (blue). (B) As in (A) except *pxb*-24xPP7 embryos were stained and PP7 smFISH probes were used. Scale bar is 50µm.

**Figure S3.**
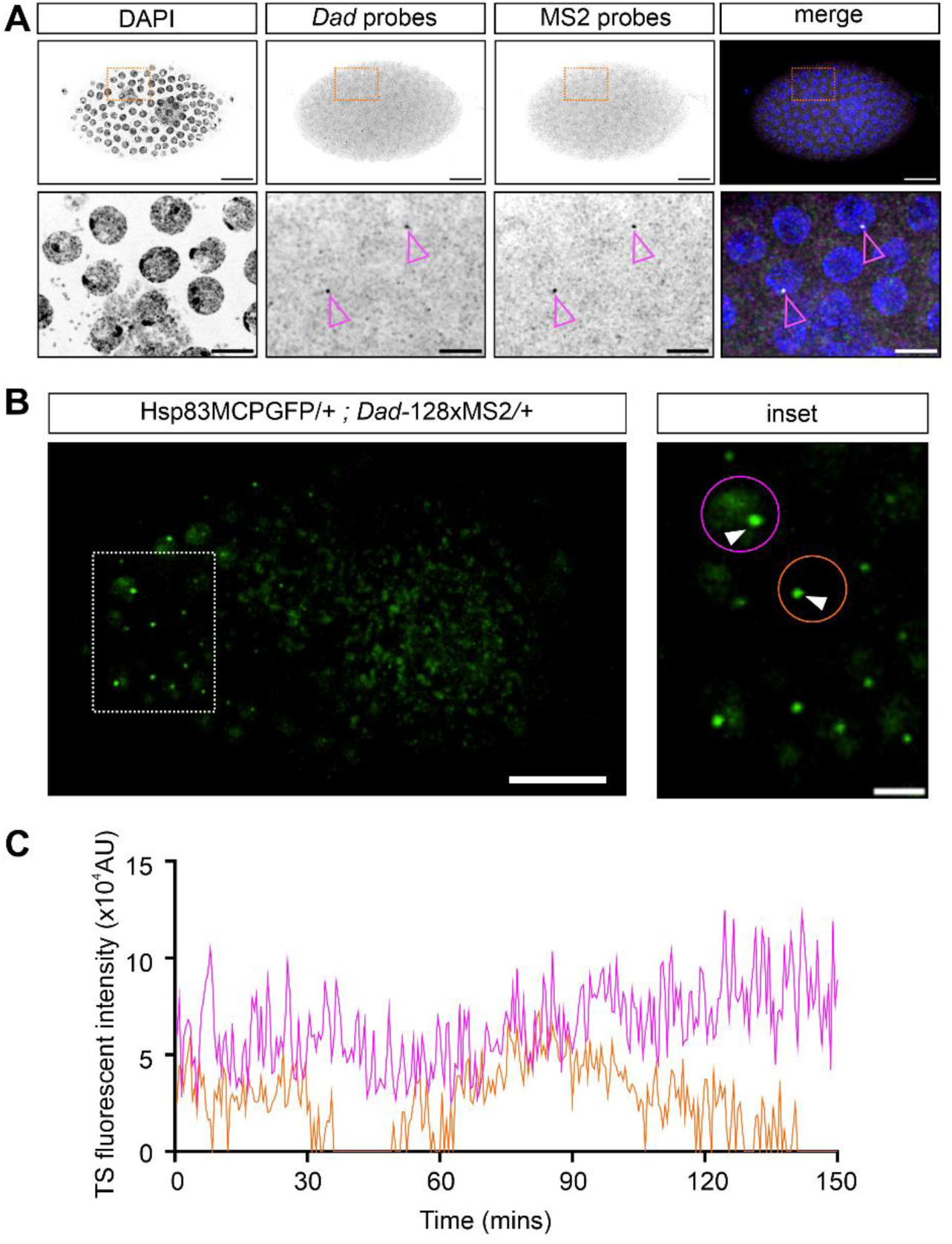
Expression of *Dad*-128xMS2 in ovarian follicle cells. (A) (Top) Confocal images of stage 8 egg chambers showing *Dad*-128xMS2 transcription in follicle cells. Nuclei are stained with DAPI and show smFISH detection of *Dad* and MS2 loops. A highlighted region of interest (orange box) is shown below. (Bottom) Higher magnification images of cells from the boxed area, TSs are marked with arrowheads. Scale bar is 20µm (top) and 5µm (bottom). (B) Still from a live imaging movie of a stage eight egg chamber from a Hsp83MCPGFP/+ ; *Dad*-128xMS2/+ female, anterior is to the left. *Dad* is transcribed in the anterior follicle cells with a region of expressing cells marked by the white box. Inset: A higher magnification image of the follicle cells highlighting two cells (purple and orange circles) with TSs (arrowheads). (C) Fluorescent intensity traces from the two cells highlighted in the inset.

**Figure S4.**
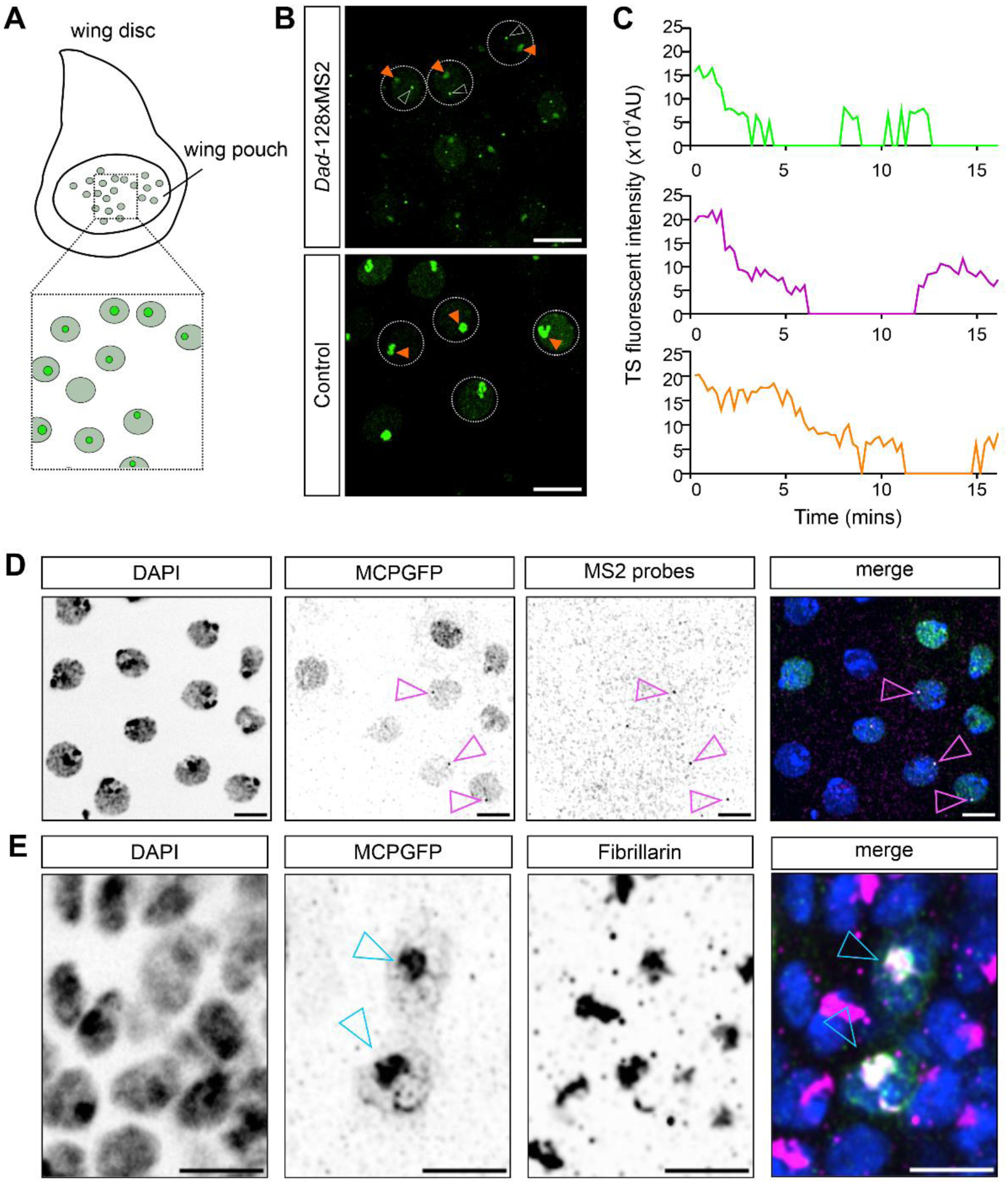
Visualising nascent transcription in the wing disc. (A) Schematic showing the third instar larval wing disc highlighting the squamous wing pouch cells that are imaged live. (B) Still from a live imaging movie of cells from Hsp83MCPeGFP/+ ; *Dad*-128xMS2/+ larvae and control cells lacking the 128xMS2 cassette. A number of cells are outlined showing active transcription sites (white arrowheads) and non-specific accumulation of MCPGFP in the nucleoli (orange arrowheads). Scale bar is 5µm. (C) TS traces from outlined cells in (B) showing transcriptional bursting. (D) Confocal images of *Dad*-128xMS2 expression in wing disc cells from Hsp83MCPeGFP ; *Dad*-128xMS2 3^rd^ instar larva. Nuclei are stained with DAPI and show MCPGFP marked Dad TSs and smFISH detection of MS2 loops. Transcription sites are marked with arrowheads. Scale bar is 5µm. (E) Hsp83MCPeGFP expressing cells of the wing disc without any MS2 loop sequences. MCPGFP accumulation colocalises with the nucleolus marked by anti-Fibrillarin immunofluorescence. Scale bar is 5µm.

**Movie 1: Live imaging of *pxb-*24xMS2 in a nuclear cycle 14 embryo.**

**Movie 2: Live imaging of *pxb-*24xPP7 in a nuclear cycle 14 embryo.**

**Movie 3: Live imaging of *pxb-*24xMS2 and *pxb-*24xPP7 in a nuclear cycle 14 embryo.**

**Movie 4: Live imaging of *Race-*24xPP7 in a nuclear cycle 14 embryo.**

**Movie 5: Live imaging of *roX1-*24xPP7 in a nuclear cycle 14 embryo.**

**Movie 6: Live imaging of *Dad*-128xMS2 in the follicle cells of stage eight egg chambers**

**Movie 7: Live imaging of *Dad*-128xMS2 in the larval brain**

**Supplementary Table 1: Table of PCR primers used in this study**

**Supplementary Table 2: Table of smFISH probes used in this study**

